# RNA-guided cell targeting with CRISPR/RfxCas13d collateral activity in human cells

**DOI:** 10.1101/2021.11.30.470032

**Authors:** Peiguo Shi, Michael R. Murphy, Alexis O. Aparicio, Jordan S. Kesner, Zhou Fang, Ziheng Chen, Aditi Trehan, Xuebing Wu

## Abstract

While single-cell sequencing has allowed rapid identification of novel cell types or states and associated RNA markers, functional studies remain challenging due to the lack of tools that are able to target specific cells based on these markers. Here we show that targeting a single marker RNA with CRISPR/RfxCas13d led to collateral transcriptome destruction in human cells, which can be harnessed to inhibit cell proliferation or to suppress cell state transition.

## RESULTS

CRISPR–Cas13 systems allow programmable targeting of RNAs^1-4^. In bacteria, upon sequence-specific recognition of viral RNAs, the non-specific RNase activity of Cas13 becomes activated and degrades both target and non-target RNAs indiscriminately. While such collateral activity has been confirmed for all Cas13 proteins tested *in vitro*^1-4^, and has been harnessed for sensitive detection of nucleic acids^5^, early investigations found no such collateral activity in eukaryotic cells^1,4,6^. Furthermore, subsequent studies reported conflicting results^7-11^, in part due to the difficulty in separating collateral activity from off-target activity caused by partial matches to the guide RNA (gRNA), as well as the lack of a proper internal control when, in principle, every cellular RNA is down-regulated by the collateral activity.

During testing of the RfxCas13d system (also known as CasRx^4^) in knocking down a DsRed reporter in human HEK293T cells, we unexpectedly found that RfxCas13d mRNA itself was also markedly down-regulated, as indicated by the loss of GFP that is encoded by the same mRNA as RfxCas13d (RfxCas13d-2A-GFP) (**Fig. 1a**). Another non-target, co-transfected BFP, was also markedly downregulated. RT-qPCR confirmed that all these mRNAs were downregulated to similar levels (20-35%) (**Fig. 1b**). Western blot confirmed that RfxCas13d protein was almost undetectable 48 hours after co-transfection with DsRed and the DsRed gRNA (**Fig. 1c**). The loss of either BFP or GFP is entirely dependent on the presence of the targeted sequence (i.e., DsRed) (**Extended Data Fig. 1**), ruling out the possibility that BFP and GFP are off-targets with partial matches to the DsRed gRNA. Taken together, these results demonstrated collateral degradation of bystander RNAs by the RfxCas13d system in human cells upon recognition of a target RNA.

**Figure 1.**
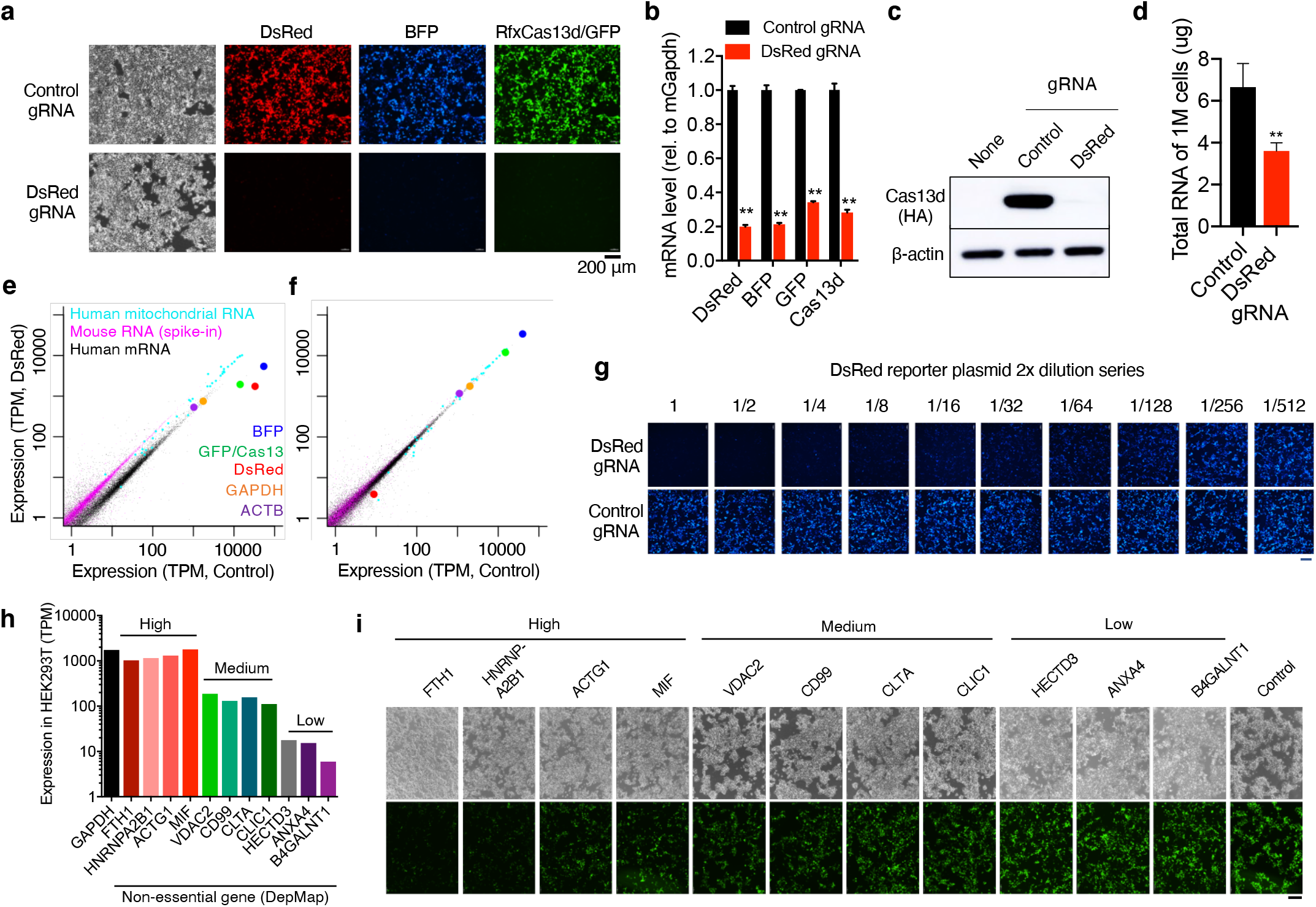
CRISPR/RfxCas13d has strong collateral activity in human cells. **a-g**, HEK293T cells were transfected with DsRed, BFP, RfxCas13d-2A-GFP, and either a control gRNA or a DsRed-targeting gRNA. Shown are fluorescence imaging (**a**), qRT-PCR (**b**), Western blot (**c**), total RNA quantified by NanoDrop (**d**), and poly(A) RNA-seq (**e-f**). For **b, d, e, f**: 5% mouse MEF-1 cells were spiked into equal number of HEK293T cells prior to RNA extraction. Mouse Gapdh (mGapdh) was used as load control in **b**. 5,000-fold less DsRed plasmid was used in **f**. Shown in **g** are BFP signal when a 2-fold dilution series of DsRed was used. **h-i**, 11 endogenous genes of various abundance (**h**) were targeted individually and the collateral degradation of RfxCas13d-2A-GFP was assayed (**i**). *: p<0.05; **: p<0.01. TPM: tags per million. **a, g, i**, scale bar, 200 μm.

Globally, the collateral activity induced by DsRed targeting led to a 46% reduction in total RNA (**Fig. 1d**), which is predominantly composed of ribosomal RNAs (rRNAs). Accordingly, we observed rRNA fragmentation and a significant reduction of RNA integrity (**Extended Data Fig. 2**). To systematically quantify the impact of RfxCas13d collateral activity on the transcriptome, we performed poly(A) RNA-seq on HEK293T cells using the DsRed-targeting RfxCas13d system in triplicate. To account for any global reduction in RNA abundance, we used an equal number of cells in each sample and spiked in 5% untransfected mouse MEF-1 cells prior to RNA extraction and sequencing. When normalized to the spike-in mouse RNAs, we observed very strong decrease of DsRed (95%), BFP (90%), and RfxCas13d/GFP (85%) RNAs (**Fig. 1e**), consistent with the strong knockdown at the protein level (**Fig. 1a/c**). Globally, almost the entire human transcriptome was down-regulated by half (median decrease of 46%), including the commonly used internal controls GAPDH and ACTB (**Fig. 1e**). This near-uniform decrease in rRNA and mRNA abundance confirmed the lack of specificity for the collateral activity of RfxCas13d, and underscored the difficulty in detecting collateral activity without a spike-in control. Interestingly, human mitochondrial RNAs are less affected by the collateral activity (**Fig. 1e**), presumably because these RNAs are shielded by the mitochondrial membrane and thus may be used as internal controls. The much stronger decrease of GFP and BFP compared to endogenous RNAs suggests that newly expressed genes may be more affected by the collateral activity of RfxCas13d. Taken together, our spike-in based RNA-seq quantification revealed the transcriptome-wide impact of RfxCas13d collateral activities in human cells.

Our observation of strong collateral RNA degradation by RfxCas13d is unexpected, given that the same RfxCas13d system was previously shown to be extremely specific in mammalian cells, with no off-target activity detected by RNA-seq when targeting two endogenous genes (ANXA4 and B4GALNT1)^4^. However, we noticed that both targets are lowly expressed (<1% of GAPDH). As each target RNA molecule can only activate a single RfxCas13d/gRNA complex, collateral damage to the transcriptome is expected to scale with the abundance of the target. Indeed, when DsRed was expressed at a level similar to ANXA4 and B4GALNT1, no systematic collateral activity was observed by RNA-seq analysis (**Fig. 1f**). Furthermore, using a serial dilution of the DsRed plasmid, we found that the collateral activity (as indicated by the loss of co-transfected non-target BFP) decreases monotonically with decreased abundance of the target (DsRed) (**Fig. 1g**).

We next targeted 11 endogenous mRNAs expressed at various levels, focusing on non-essential genes (DepMap^12^) (**Fig. 1h**). Targeting four highly expressed genes whose expressions are comparable to GAPDH led to a marked decrease of RfxCas13d/GFP (i.e., strong collateral activity) (**Fig. 1i**). In contrast, targeting three lowly expressed genes that are at ∼1% of GAPDH, including ANXA4 and B4GALNT1 tested in the original RfxCas13d study^4^, showed no apparent decrease of GFP (**Fig. 1i**). Targeting four medium abundance genes with expression about 10% of GAPDH resulted in moderate loss of GFP (**Fig. 1i**). These results show that targeting endogenous RNAs can also result in collateral activity, and that the abundance of the target RNA strongly influences the extent of this collateral activity.

In bacteria, CRISPR/Cas13-mediated destruction of the transcriptome is known to result in host dormancy when induced by viral infection^13^. Similarly, we observed that collateral activity induced by targeting of DsRed strongly affects cell viability and proliferation (as measured by WST-1 assay, **Fig. 2a**) in a target (i.e., DsRed) abundance dependent manner (**Extended Data Fig. 3**) without causing significant cell death or apoptosis (**Extended Data Fig. 4**). Compared to controls, targeted cells showed a marked decrease in DNA replication activity (reduced EdU incorporation, **Fig. 2b**). Intriguingly, unlike the nuclei in control cells that have a smooth outline and more uniformly distributed chromatin, the nuclei of DsRed-targeted cells often have a rugged and irregular boundary and contain dense chromatin clumps (**Fig. 2c**). This chromatin collapse phenotype is similar to that of cells treated with RNase^14,15^, consistent with activated RfxCas13d functioning as a non-specific RNase. In addition to impairing DNA replication, the collapse of chromatin may also shutdown global transcription, which potentially underlies the heightened impact on newly expressed genes. Targeting the 4 abundant but non-essential endogenous mRNAs mentioned previously led to a similar reduction in cell proliferation (**Fig. 2d**), whereas targeting mRNAs of medium abundance resulted in a moderate decrease in cell viability and no significant change for low-abundance targets (**Fig. 2d**).

**Figure 2.**
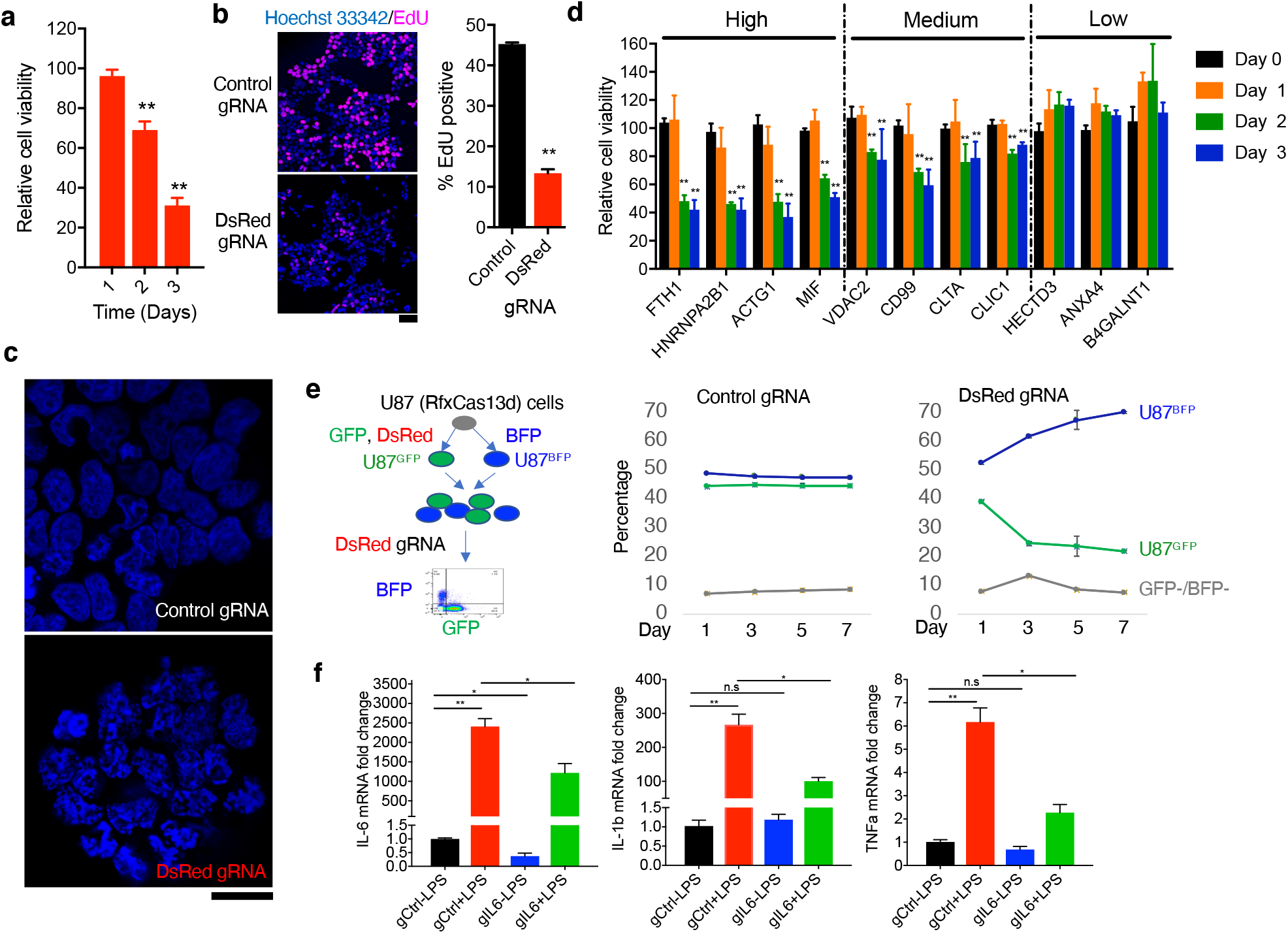
Harnessing CRISPR/RfxCas13d collateral activity for RNA-guided cell targeting. **a-c**, HEK293T cells were transfected with RfxCas13d, DsRed, and either a DsRed-targeting gRNA or a non-targeting control gRNA. Shown are (**a**) cell viability / proliferation (relative to control gRNA) as measured by WST-1 assay, (**b**) EdU incorporation assay, and (**c**) nuclear morphology with DAPI staining. **d**, WST-1 assay for targeting endogenous non-essential mRNAs. **e**, Selective depletion of DsRed-expressing U87^GFP^ cells from a mixture of U87^GFP^ and U87^BFP^ cells by using RfxCas13d (stably expressed) and a DsRed gRNA (transfected on day 0). **f**, Expression of IL-6, IL-1b, and TNFa mRNAs in human PBMCs with and without LPS treatment. Cells express RfxCas13d and either an IL-6 targeting gRNA (gIL6) or a control gRNA (gCtrl). *: p<0.05; **: p<0.01. n=3 for all panels. Scale bar, 50 μm (**b**,**c**).

To further demonstrate the utility of RfxCas13d in selectively depleting a sub-population of cells (such as cancer cells) expressing a specific marker RNA that has no functional role in the cell, we mixed two human U87 glioblastoma cell lines stably expressing either GFP (U87^GFP^) or BFP (U87^BFP^), and then expressed RfxCas13d and a gRNA targeting DsRed in the cell mixture. The target DsRed is only expressed in U87^GFP^ cells (**Fig. 2e**). In this competitive growth assay, the fraction of the targeted U87^GFP^ cells decreased over time, whereas U87^BFP^ cells increased (**Fig. 2e**). These results demonstrated that the collateral activity of RfxCas13d can be harnessed for sequence-specific inhibition of cell proliferation, regardless of whether or not the target RNA plays any functional role in the cell.

The global down-regulation of the transcriptome (**Fig. 1e**), and in particular newly expressed genes (BFP and GFP), suggests that one can target a single upregulated RNA with RfxCas13d to suppress a concerted transcriptional activation program, which in turn blocks cell state transition. One such example is the simultaneous upregulation of multiple cytokines during inflammation. For instance, SARS-CoV-2 infection can lead to upregulation of TNFα, IL-1β, IL-6, and other cytokines among immune cells in the lung, which activates a positive-feedback loop leading to hyperinflammation and resulting in organ and tissue damage (cytokine storm)^16,17^. Using lipopolysaccharide (LPS)-based simultaneous induction of IL-6, IL-1β, and TNFα in human peripheral blood mononuclear cells (PBMCs) as a model system, we show that with a single gRNA targeting IL-6, RfxCas13d down-regulated not only IL-6, but also IL-1β and TNF*α* (**Fig. 2f**). Similar results were observed in the human monocyte cell line U937 (**Extended Data Fig. 5**). In both PBMCs and U937 cells, the collateral down-regulation of IL-1β and TNF*α* depends on IL-6 upregulation by LPS, as IL-1β and TNF*α* are not affected in cells that have not been treated with LPS (thus low IL-6 expression), consistent with target abundance dependent activation of RfxCas13d collateral activities. Importantly, IL-6 knockdown by siRNA has no effect on IL-1β or TNF*α* expression, with or without LPS treatment (**Extended Data Fig. 6**), strongly suggesting that IL-6 induced RfxCas13d collateral activity, but not IL-6 down-regulation *per se*, causes the down-regulation of IL-1β and TNF*α*. IL-6 targeting by RfxCas13d did not cause more cell death than a control gRNA (**Extended Data Fig. 7**), thus allowing the suppression of the transition to a pro-inflammatory state without killing the targeted cells.

In summary, we show that the CRISPR/RfxCas13d system has strong collateral activity in human cells and can be harnessed to inhibit the proliferation of specific cell types or to suppress the transition of cell states based on marker RNAs. While these results call for caution in using CRISPR/RfxCas13d for targeted RNA knockdown, we propose that further development of such a programmable RNA-guided cell targeting platform has the potential to transform functional studies of the myriad cell types and cell states uncovered by single-cell sequencing technologies. It may also advance therapeutic development by allowing sequence-specific targeting of cancer cell proliferation, immune cell inflammatory state, and other pathogenic cell types and cell states.

## Data availability

RNA-seq data were deposited in Gene Expression Omnibus (GEO) with the accession number GSE155134.

## Acknowledgements

We thank Hanrui Zhang, Muredach Reilly, and Peter Sims for generously sharing cell lines and members of the laboratory of X.W. and the Cardiometabolic Genomics Program for advice and discussions. M.R.M. is supported by an American Heart Association postdoctoral fellowship. X.W. is supported by NIH grant 1DP2GM140977 and Pershing Square Sohn Cancer Research Alliance. This research was funded in part through the NIH/NCI Cancer Center Support Grant P30CA013696 and used the Genomics and High Throughput Screening Shared Resource and CCTI Flow Cytometry Core. The CCTI Flow Cytometry Core is supported in part by the Office of the Director, National Institutes of Health under awards S10RR027050 and S10OD020056. The content is solely the responsibility of the authors and does not necessarily represent the official views of the National Institutes of Health.

## Author contributions

P.S., M.R.M., and X.W. conceived the project. P.S. performed all experiments and data analysis with assistance from M.R.M., A.O.A., Z. F., Z. C., A. T., and J. S. K. P.S. and X.W. drafted the manuscript with input from all authors.

## METHODS

### Cell culture

HEK293T, MEF-1, U87, U937 cell lines were used in this research. HEK293T was purchased from ATCC. MEF-1, U87, and U937 cell lines were gifts from Muredach P. Reilly, Peter A. Sims. and Hanrui Zhang, respectively. HEK293T, MEF-1 and U87 cells were cultured in DMEM with 4.5 g/L D-Glucose, supplemented with 10% fetal bovine serum (FBS), no antibiotic was added. Cells were passaged upon reaching 80%–90% confluency. U937 was cultured in RPMI1640 with 10% FBS, and without antibiotics. U937 cell density was maintained between 1 × 10^5 and 2 × 10^6 cells/ml. All cells were cultured at 5% CO_2_ and 37 °C. All cells were routinely tested for mycoplasma contamination using MycoAlert™ Mycoplasma Detection Kit (Lonza, LT07-418).

### Plasmids

RfxCas13d-2A-GFP was expressed using the plasmid pXR001: EF1a-CasRx-2A-EGFP (Addgene #109049, ref^4^). In the competitive growth assay described in Fig. 2e, GFP was replaced with puroR. Guide RNAs (gRNA), except gIL6, were cloned into the plasmid RfxCas13d gRNA cloning backbone (Addgene #109053, ref^4^) using oligos listed in Supplementary Table 1. gIL6 was cloned into the all-in-one plasmid hU6-DR_BsmBI-EFS-RfxCas13d-NLS-2A-Puro-WPRE (Addgene #138147, ref^18^). DsRed plasmid was modified from pLenti-DsRed_IRES_EGFP (Addgene #92194, ref^19^) by deleting IRES-EGFP. BFP was expressed with a plasmid originally used for the CRISPRi gRNA expression (Addgene # 62217, ref^20^).

### Plasmid transfection

All plasmids were prepared using the NuceloSpin Plasmid Transfection-grade kit (MACHEREY-NAGEL, 740490.250). Plasmids were transfected with Lipofectamine 3000 (Invitrogen, L3000015) according to the manufacturer’s protocol. Briefly, HEK293T cells were seeded at a density of 6 × 10^5 per well in a 6-well plate and transfected the next day. Opti-MEM I Reduced-Serum Medium (Gibco, 31985062) was prewarmed at 37 °C water bath for 30 min prior to transfection. For each well, 7.5 μl lipofectamine 3000 reagent was mixed with 125 μl Opti-MEM I Reduced-Serum Medium, then mixed with 2500 ng DNA in 125 μl Opti-MEM I Reduced-Serum Medium. After 15 min at room temperature, the mix was added to cells. Media was not changed after transfection. Cells were harvested 48 hours after transfection for gene expression analysis, western blot, flow cytometry, and RT-qPCR assays.

### siRNA transfection

A non-targeting control siRNA pool (Horizon Discovery, D-001810-10-05) and a human IL-6 siRNA pool (Horizon Discovery, LQ-007993-00-0005) were transfected using Lipofectamine™ RNAiMAX Transfection Reagent (Invitrogen, 13778) according to the manufacturer’s protocol. PBMCs were seeded at a density of 1 × 10^6 per well in a 6-well plate. Opti-MEM I Reduced-Serum Medium (Gibco, 31985062) was prewarmed at 37 °C water bath for 30 min prior to transfection. For each well, 7.5 μl lipofectamine RNAiMAX reagent was mixed with 125 μl Opti-MEM I Reduced-Serum Medium and then mixed with 25 pmol siRNA in 125 μl Opti-MEM I Reduced-Serum Medium. After 5 min at room temperature the mix was added to cells. Media was not changed after transfection. PBMCs were treated with 1 μg/ml LPS for 8 hours 48 hours after transfection, then cells were collected for RNA extraction and MEF-1 spike-in RT-qPCR.

### Quantitative reverse transcription PCR (RT-qPCR)

Viable treated cells were counted and mixed with 5% (by number) untransfected mouse MEF-1 cells. Total RNA was isolated using the RNA isolation kit (MACHEREY-NAGEL, 740984.250). Synthesis of cDNA was done using the SuperScript™ IV Reverse Transcriptase (Invitrogen, 18090050). Quantitative RT–qPCR was performed with Real-Time PCR System (Applied Biosystems, Quantstudio 7 Flex), using SYBR Green Master Mix (Applied Biosystems, A25741). Relative transcript abundance was normalized to mouse *Gapdh (mGapdh)*. Primer sequences can be found in Supplementary Table 2.

### Poly(A) RNA-seq

HEK293T cells were seeded at a density of 6 × 10^5 per well in a 6-well plate. 2 × 10^5 MEF-1 cells were seeded in a 6-well plate. For the experiment described in Fig. 1e, 625 ng of each plasmid (RfxCas13d-2A-GFP, DsRed, BFP, and control or DsRed targeting gRNA) were transfected into HEK293T cells according to the above plasmid transient transfection protocol (2500 ng DNA total per well). For Fig. 1f, 5,000-fold less DsRed plasmid was used (0.125 ng). Cells were harvested 48 hours after transfection. For each sample, 5 × 10^4 MEF-1 cells were spiked into 1 × 10^6 viable HEK293T cells, followed by RNA extraction, poly(A) RNA pulldown, and sequencing library construction using Illumina TruSeq chemistry. Libraries were then sequenced using Illumina NovaSeq 6000 at Columbia Genome Center. RTA (Illumina) was used for base calling and bcl2fastq2 (version 2.19) for converting BCL to fastq format, coupled with adaptor trimming. kallisto (0.44.0) was used to perform a pseudoalignment to indices created from human and mouse transcriptomes (Human: GRCh38; Mouse: GRCm38). Gene expression in TPM (tags per million) were quantified using kallisto. TPM values were then normalized to the total counts of mouse RNAs in each sample.

### Western Blot

HEK293T cells were seeded at a density of 6 × 10^5 per well in a 6-well plate. A total of 2500 ng DNA (625 ng RfxCas13d-2A-GFP, 625 ng DsRed, 625 ng BFP, and 625 ng control or DsRed targeting gRNA plasmids) were transfected into HEK293T cells according to the above plasmid transient transfection protocol. Cells were harvested 48 hours after transfection. Cells were gently washed by cold PBS and lysed with RIPA buffer (Sigma, R0278) supplemented with protease inhibitors (Roche, 4693132001). Cell lysates were cleared by centrifugation for 15 min at 12,000 x g, 4°C. Protein lysates were loaded with LDS Sample Buffer (Invitrogen, NP0007) and Reducing Agent (Invitrogen, NP0004) after heating at 70°C for 10 min. Protein were separated by SDS - PAGE and subsequently transferred to PVDF membrane. Membranes were blocked in PBS with 5% non-fat milk and 0.1% Tween-20 for 1 h at room temperature and probed with appropriate primary antibodies overnight at 4°C. Primary antibodies used: HA (Sigma, H9658, 1:1000), β-Actin (Sigma, A5441, 1:2000), LC3 (Proteintech, 14600-1-AP, 1:1000). HRP conjugated secondary antibodies were incubated for 1 h at room temperature. Blots were imaged using West Pico PLUS Chemiluminescent Substrate (Thermo fisher, 34580).

### Cell proliferation assays (EdU assays)

HEK293T cells were seeded at a density of 3 × 10^5 per well in a 12-well plate with Poly-D-Lysine coated German glass coverslips (Neuvitro, H-15-PDL). A total of 1,000 ng plasmids were transfected for 48 hours, and then cells were incubated with EdU for 4 hours, cells were subsequently fixed by 4% paraformaldehyde (PFA). EdU assays were completed with Click-iT Plus EdU Cell Proliferation Kit (Invitrogen, C10640) according to the manufacturer’s protocol.

### Cell viability assays (WST-1)

HEK293T cells were seeded at a density of 1.8 × 10^4 per well in a 48-well plate. Cells were transfected with 250 ng DNA per well for the indicated time of figures. Cells were incubated with the WST-1 reagent (Sigma, 11644807001) for 2 hours. The absorbance of the samples was measured using a microplate reader at 440 nm.

### Apoptosis assays

Annexin V and DAPI staining assays were performed using the apoptosis detection Kit (BioLegend, 640930) according to the manufacturer’s protocol. Briefly, 48 hours after transfection, cells were washed twice with cold cell staining buffer (Biolegend, 420201) and resuspended in binding buffer. APC-Annexin V and DAPI (Alternative for 7-AAD of the kit) were used for staining. Cells were incubated for 15 min at room temperature and analyzed by flow cytometry. Data were analyzed using FCS Express 7.10.

For cleaved Caspase 3 staining assays, HEK293T cells were seeded at a density of 3 × 10^5 per well in a 12-well plate with Poly-D-Lysine coated German glass coverslips. Cells were fixed by 4% PFA 48 hours after transfection. Cleaved Caspase 3 antibody (Cell signaling technology, 9661T, 1:500) was used for detecting the expression of cleaved Caspase 3. Confocal images were captured with an LSM T-PMT confocal laser-scanning microscope (Carl Zeiss).

### Lentivirus and stable cell line generation

We generated lentivirus with pCMV-dR8.91 and pMD2.G packaging system (gifts from Jonathan Weissman). Briefly, packaging vectors were transfected into HEK293T cells using Lipofectamine 3000 according to the manufacturer’s protocol. Lentiviruses were harvested 48 hours post transfection, aliquoted and stored at –80°C for future use. For U87 or U937 stable cell lines generation, cells were seeded in a 6-well plate and infected by lentivirus with 10 μg/ml of polybrene for 48 hours. Stable cells were then isolated by antibiotic selection. RfxCas13d RNA expression levels were quantified by RT-qPCR to ensure that samples had comparable expression levels. All experiments involving stable cell lines were performed with low passage cells.

### Competitive growth assay

A human U87 glioblastoma cell line stably expressing RfxCas13d (no GFP fusion) was generated using lentiviral integration followed by puromycin selection. Two cell lines were subsequently derived using lentiviral integration of either GFP or BFP followed by FACS. In the absence of treatment, the two cell lines, termed U87^GFP^ cells and U87^BFP^ cells respectively, have similar doubling time. The U87^GFP^ cell line additionally expresses the DsRed reporter. The two cell lines were mixed at 1:1 ratio on day 0, then an in vitro transcribed gRNA targeting DsRed was transfected using Lipofectamine 3000 according to the manufacturer’s protocol. Mock transfection and transfection of a non-targeting gRNA were used as control. The fraction of U87^GFP^ and U87^BFP^ cells were determined using flow cytometry by detecting the GFP and BFP expression.

### Human peripheral blood mononuclear cells (PBMCs) isolation and spinfection

Human blood samples were obtained from New York Blood Center. PBMCs were isolated by Ficoll density gradient centrifugation (Sigma, GE17-5442-02). To introduce RfxCas13d and IL6-gRNA into PBMCs, lentivirus spinfection method was performed. Briefly, 4 × 10^6 PBMCs were seeded in a 6-well plate with 1 ml medium and 1 ml lentivirus that generated by the above lentivirus generation protocol was added into wells, with the final concentration of polybrene at 10 μg/ml. The plate was centrifuged for 90 min at 950 g, 32 °C, and the medium was changed after 24 hours of incubation. No selection was used. Cells were harvested after 8 h of 1 μg/ml LPS treatment, MEF-1 spike-in RT-qPCR was subsequently performed to detect the expression of cytokines. The expression of RfxCas13d RNA was quantified by RT-qPCR to ensure that samples have comparable expression levels.

### Statistical analyses

In general, at least three biological replicates were performed for each experiment. All statistical analyses were performed using the GraphPad Prism software 9.0. Student’s t-test was used for comparison between two groups. Results in graphs are expressed as mean ± SD or mean ± SEM.

---

**Supplementary Table 1:**
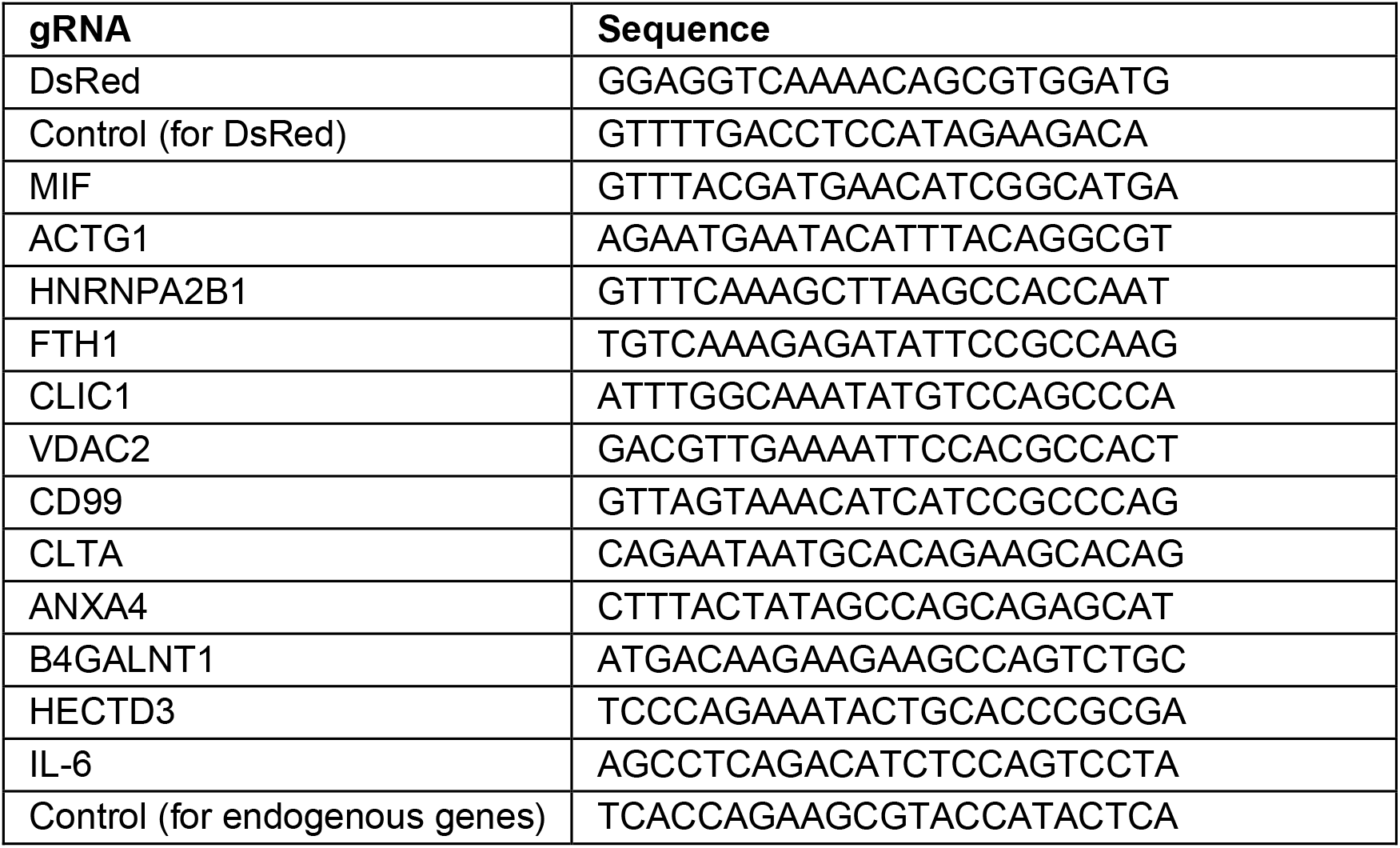
Guide RNA (gRNA) spacer sequences.

**Supplementary Table 2:**
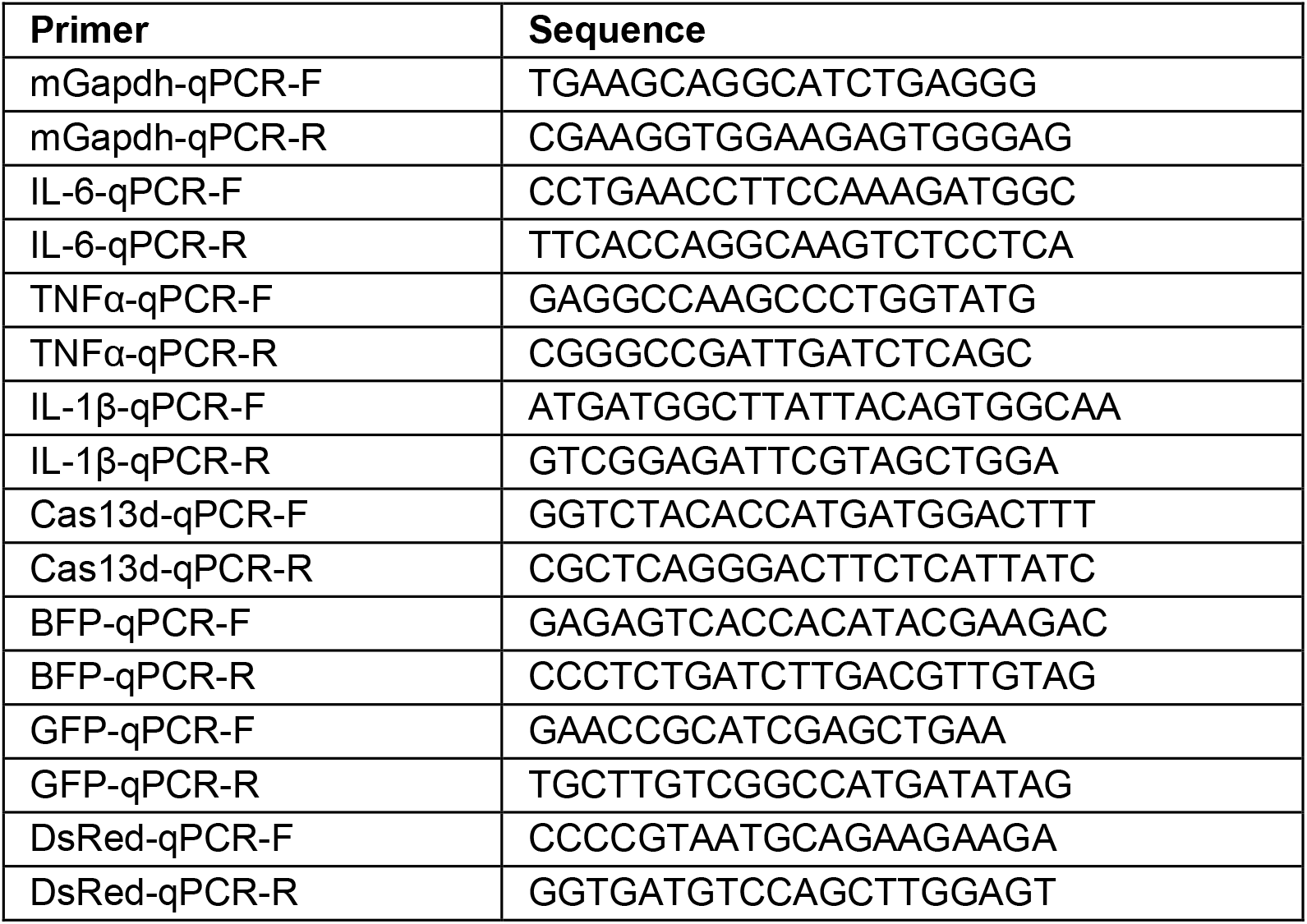
RT-qPCR primers sequences.

**Extended Data Figure 1:**
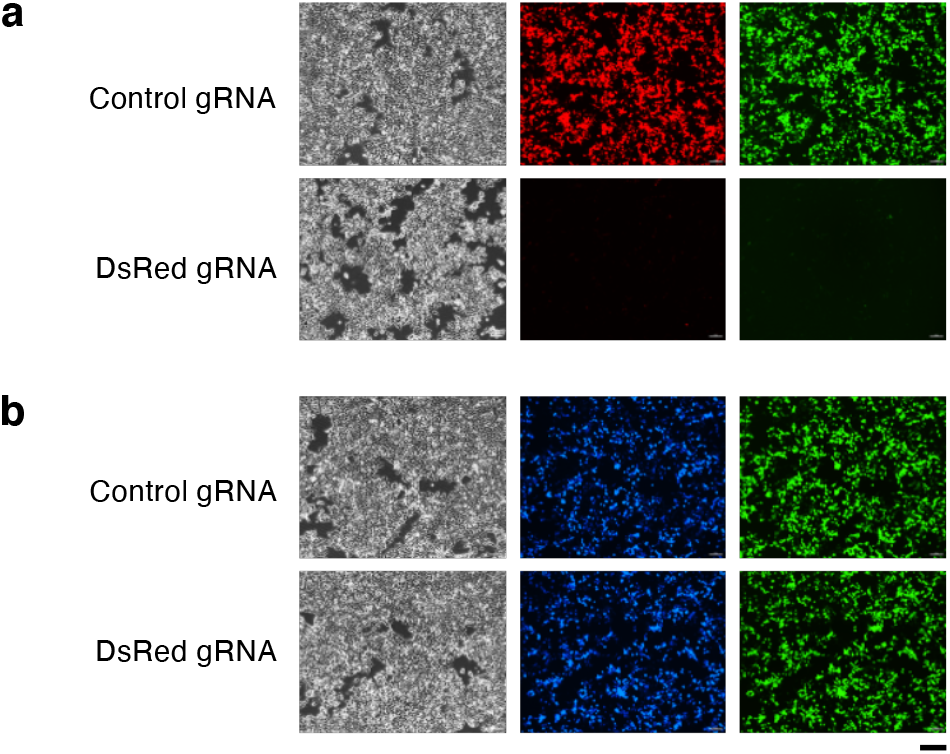
The loss of BFP/GFP depends on the target of the gRNA. (i.e., DsRed). **a**, Cells were transfected with RfxCas13d-2A-GFP, DsRed, and either a DsRed targeting gRNA or a non-targeting control gRNA. **b**, same as **a** but replacing DsRed with BFP. Note that BFP/GFP are only down-regulated when DsRed is present and targeted by RfxCas13d loaded with a DsRed gRNA. Scale bar, 200 μm.

**Extended Data Figure 2:**
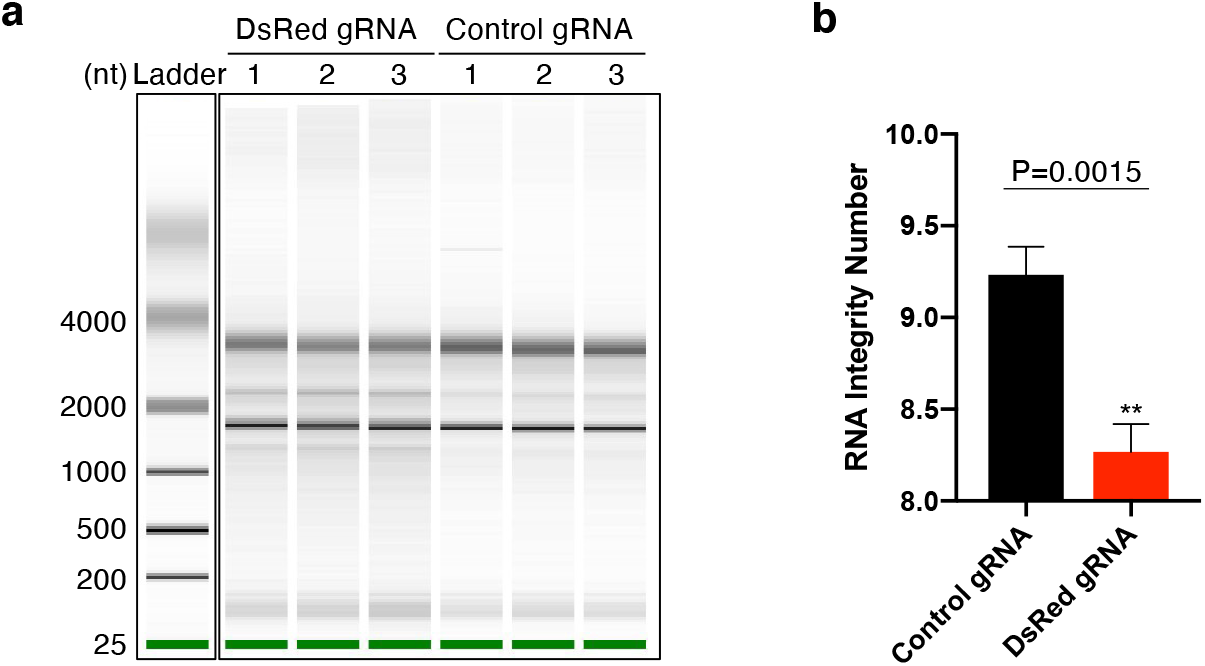
Targeting DsRed with CRISPR/RfxCas13d led to rRNA degradation (**a**, BioAnalyzer) and a significant reduction of RNA integrity number (**b**). N=3.

**Extended Data Figure 3:**
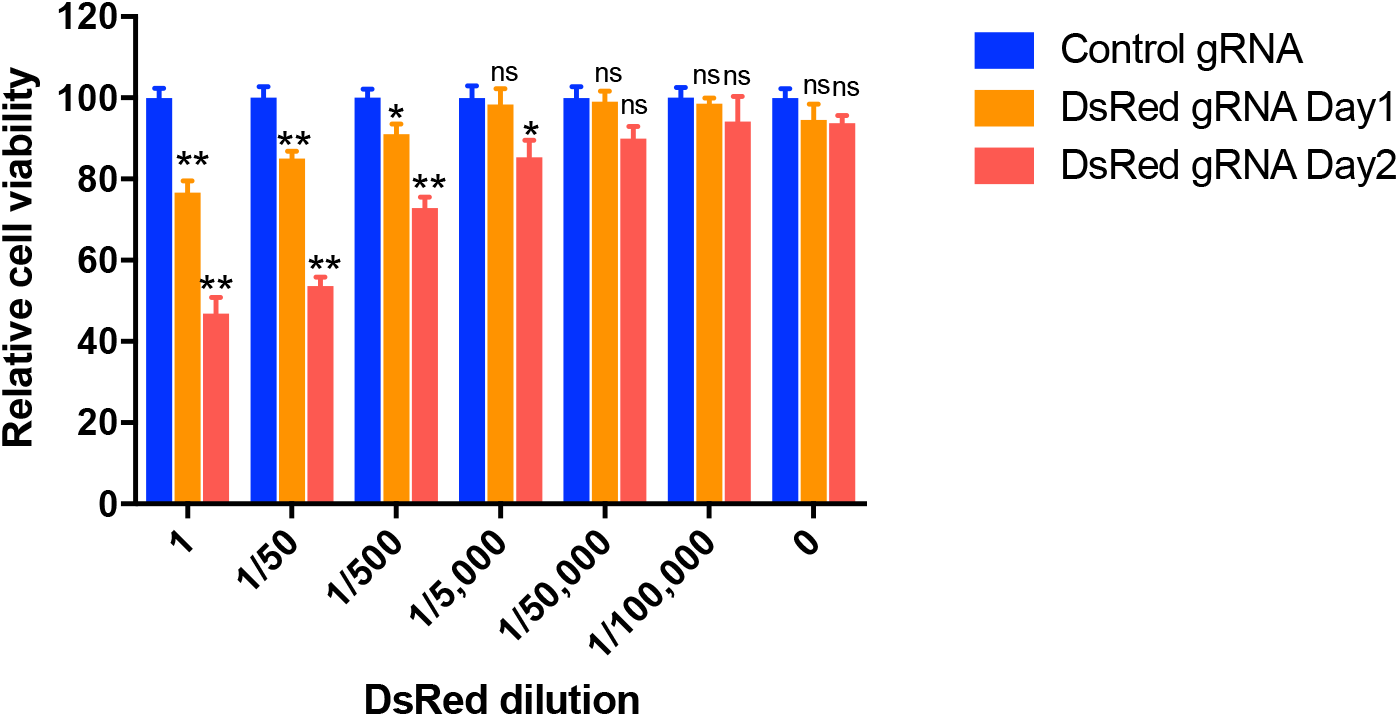
Change in cell viability/proliferation (WST-1 assay) in cells transfected with varying amount of DsRed plasmids, together with a fixed amount of RfxCas13d and gRNA plasmids.

**Extended Data Figure 4.**
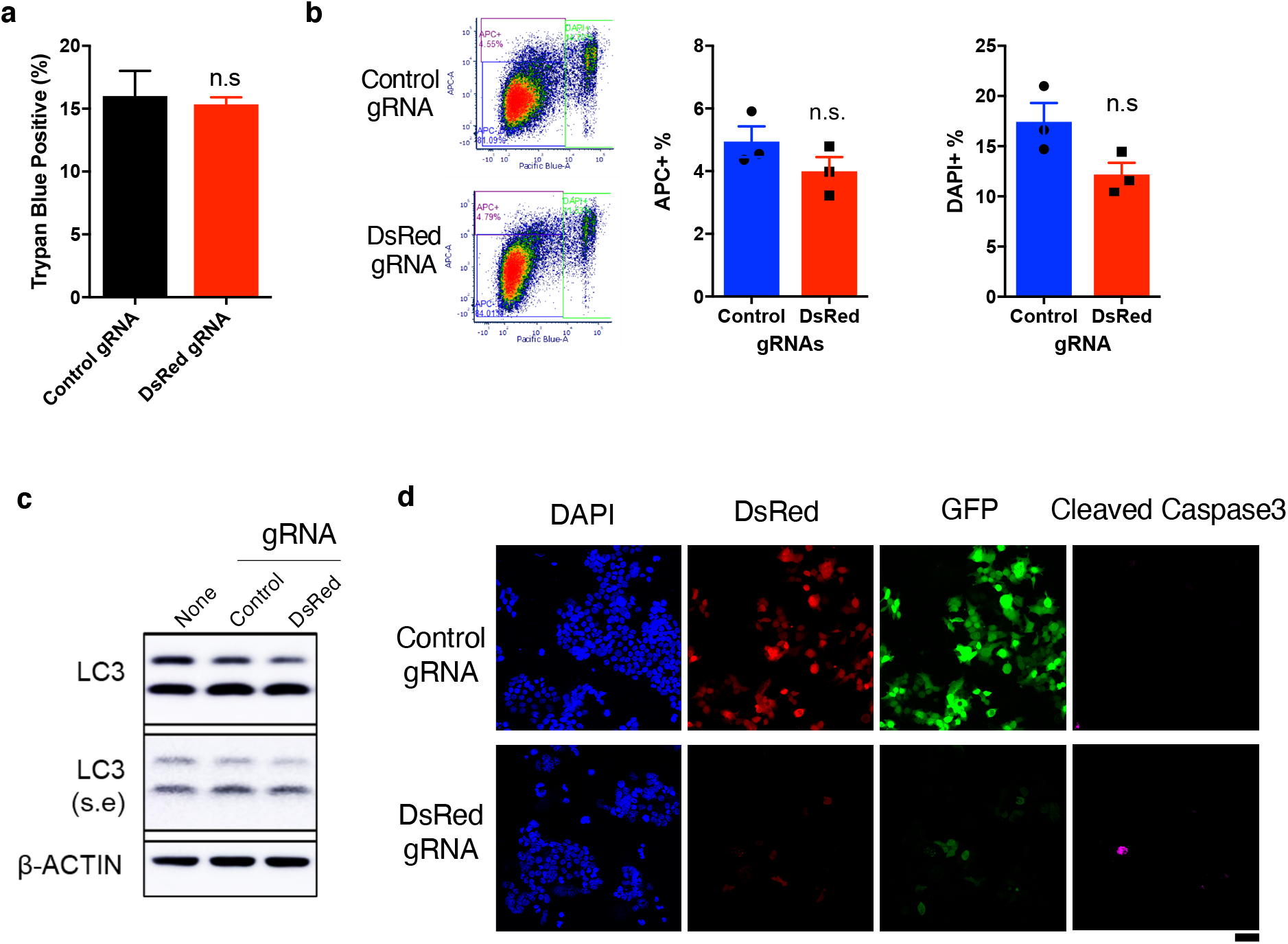
DsRed-induced RfxCas13d collateral activity does not cause cell death. HEK293T cells were transfected with DsRed, RfxCas13d-2A-GFP, and either a DsRed-targeting gRNA or a non-targeting control gRNA. No difference was detected with trypan blue-based cell viability assay (**a**), Annexin V APC / DAPI cell apoptosis assay (**b**), LC3 cleavage (autophagy) (**c**), or Caspase3 cleavage (apoptosis) (**d**). Scale bar, 50 μm.

**Extended Data Figure 5.**
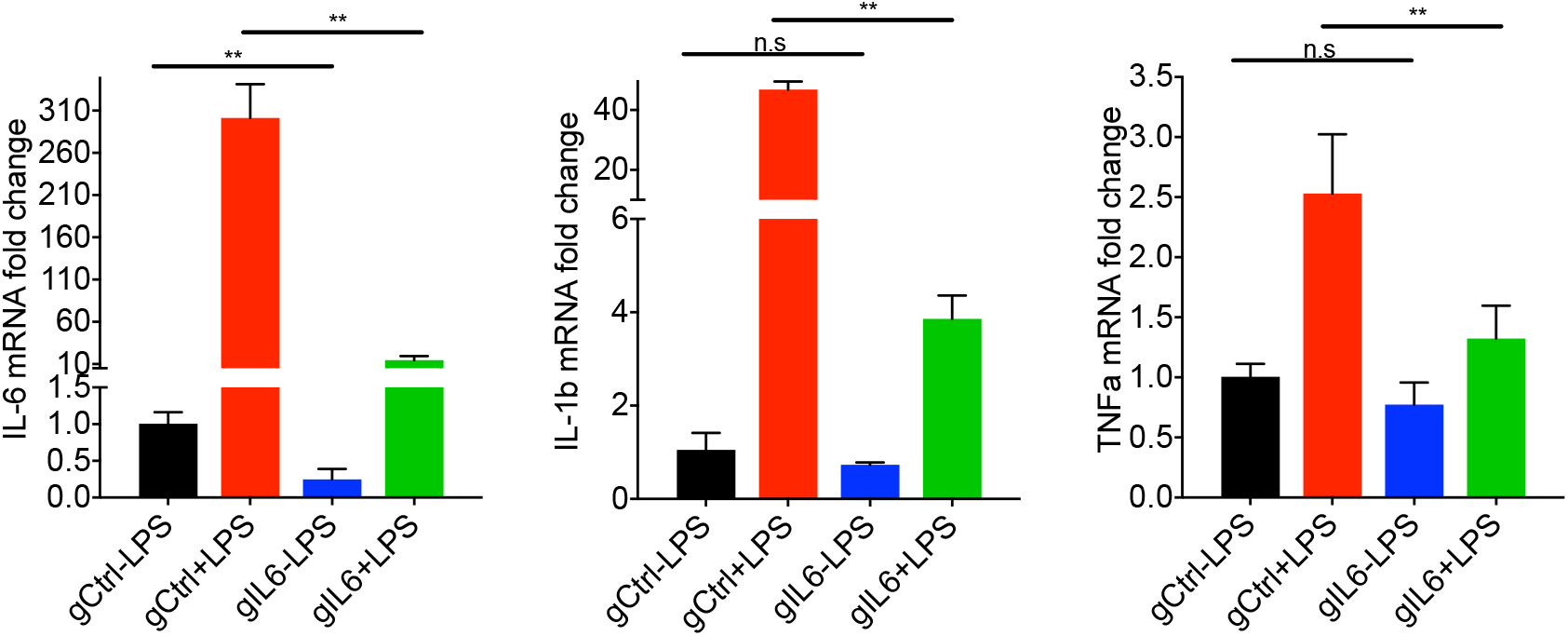
Inhibiting LPS-activation of IL-6 by RfxCas13d also inhibits the activation of IL-1β and TNF*α* in U937 human monocyte cells. Same as Fig. 2f but in U937 cells.

**Extended Data Figure 6.**
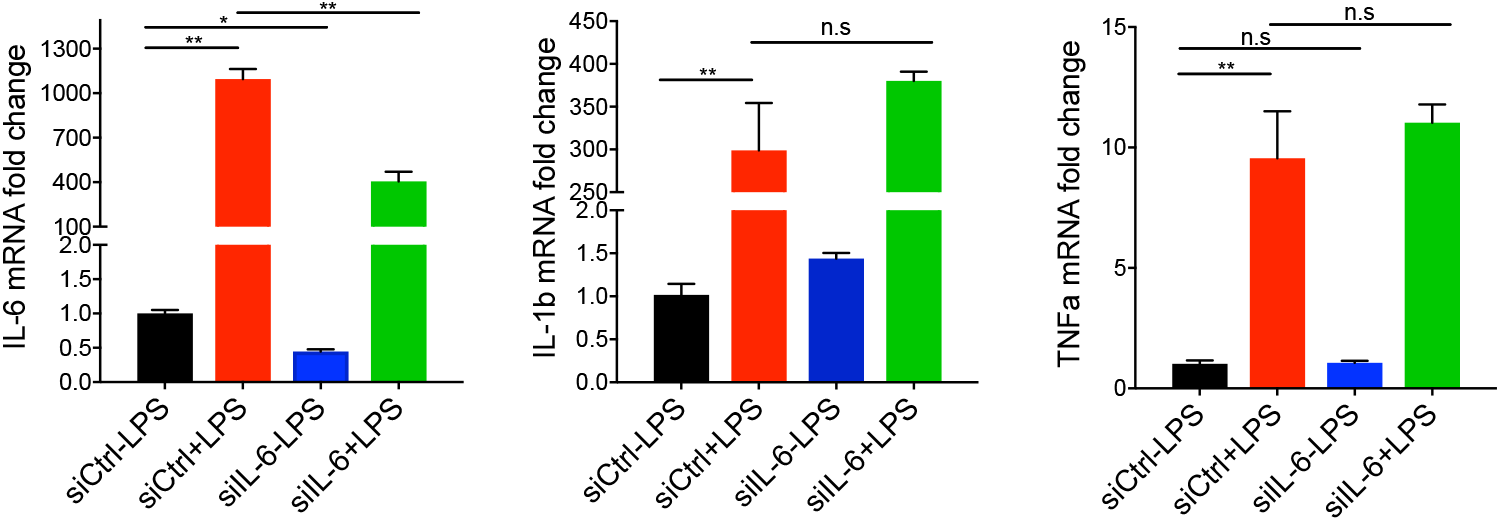
IL-6 knockdown by siRNA has no effect on IL-1β or TNF*α* expression. Human PBMCs were transfected with either an IL-6 targeting siRNA (siIL-6) or a control siRNA (siCtrl). LPS was added 48 hours later. The expression of IL-6, IL-1b, and TNFa were measured by RT-qPCR 8 hours after LPS treatment.

**Extended Data Figure 7.**
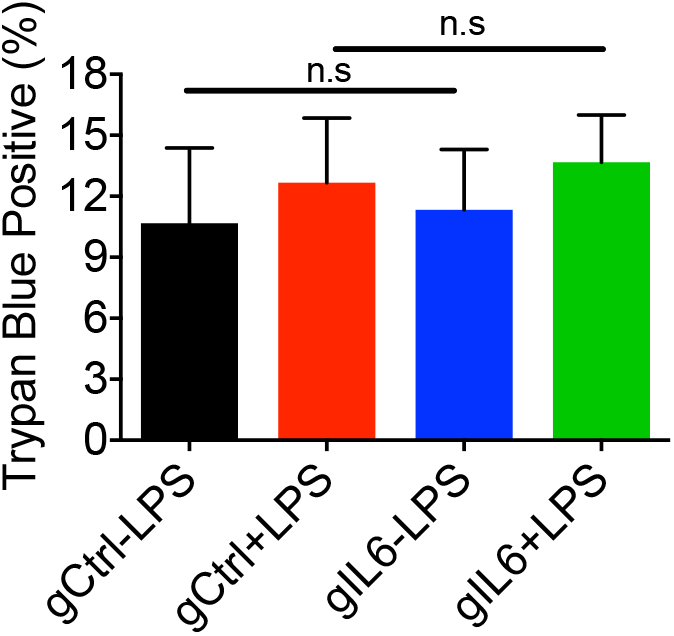
RfxCas13d targeting of IL-6 has no significant impact on cell viability in PBMC. N=3.

